# Extracellular Matrix Instability and Chronic Inflammation Underlie Maladaptive Right Ventricular Pressure Overload Remodeling and Failure in Male Mice

**DOI:** 10.1101/2024.04.03.588013

**Authors:** Ilaria Russo, Wen Dun, Swasti Mehta, Sowda Ahmed, Christos Tzimas, Nobuaki Fukuma, Emily J. Tsai

## Abstract

**Background:** Right ventricular dysfunction (RVD) portends increased death risk for heart failure (HF) and pulmonary arterial hypertension (PAH) patients, regardless of left ventricular function or etiology. In both, RVD arises from the chronic RV pressure overload, and represents advanced cardiopulmonary disease. RV remodeling responses and survival rates of HF and PAH patients, however, differ by sex. Men develop more severe RVD and die at younger ages than do women. Mechanistic details of this sexual dimorphism in RV remodeling are incompletely understood. We sought to elucidate the cardiac pathophysiology underlying the sex-specific RV remodeling phenotypes, RV failure (RVF) versus compensated RVD.

**Methods:** We subjected male (M-) and female (F-) adult mice to moderate pulmonary artery banding (PAB) for 9wks. Mice underwent serial echocardiography, cardiac MRI, RV pressure-volume loop recordings, histologic and molecular analyses.

**Results:** M-PAB developed severe RVD with RVF, increased RV collagen deposition and degradation, extracellular matrix (ECM) instability, and activation and recruitment of macrophages. Despite the same severity and chronicity of RV pressure overload, F-PAB had more stable ECM, lacked chronic inflammation, and developed mild RVD without RVF.

**Conclusions:** ECM destabilization and chronic activation of recruited macrophages are associated with maladaptive RV remodeling and RVF in male PAB mice. Adaptive RV remodeling of female PAB mice lacked these histopathologic changes. Our findings suggest that these two pathophysiologic processes likely contribute to the sexual dimorphism of RV pressure overload remodeling. Further mechanistic studies are needed to assess their pathogenic roles and potential as targets for RVD therapy and RVF prevention.

**CLINICAL PERSPECTIVE:** *What is new?:*

- In a mouse model of pure PH, males but not females showed an association between ECM instability, chronic inflammation with activation of recruited macrophages, and severe RV dysfunction and failure.

*What are the clinical implications?:*

- In male HF and PH patients, enhancing ECM stability and countering the recruitment and activation of macrophages may help preserve RV function such that RVF can be prevented or delayed. Further preclinical mechanistic studies are needed to assess the therapeutic potential of such approaches.

**RESEARCH PERSPECTIVE:** *What new question does this study raise? What question should be addressed next?:*

- What mechanisms regulate RV ECM stability and macrophage recruitment and activation in response to chronic RV pressure overload? Are these regulatory mechanisms dependent upon or independent of sex hormone signaling?

## INTRODUCTION

Sex differences have been reported in the risk factors, incidence, and clinical outcomes for various cardiovascular diseases, including heart failure (HF).^1^ This sexual dimorphism, whereby men are disadvantaged relative to women, is accentuated in right ventricular remodeling and dysfunction (RVD). In male patients, the RV commonly responds to chronic pressure overload (i.e. pulmonary hypertension, PH) with maladaptive remodeling; RV contractility cannot augment sufficiently to counter the increased RV afterload. Severe RVD subsequently ensues and then devolves into RV failure (RVF). In female patients, however, the chronically, pressure-overloaded RV remodels adaptively, allowing for preservation of RV stroke work efficiency and a prolonged stage of compensated RVD.^2^ It is this observed sex difference in RV pressure overload remodeling that drives the different clinical courses of male and female patients with HF and PH. Given that RVD independently predicts death in HF and PH, there is an urgent need to understand the molecular pathophysiology of maladaptive RV remodeling in males and adaptive RV remodeling in females.

Current understanding of sexual dimorphism in RV pressure overload remodeling is largely based upon pre-clinical studies demonstrating the cardioprotection of the potent estrogen 17β-estradiol (E2) against various cardiac stressors.^3–9^ The mechanistic details of E2 cardioprotection in the RV, however, are not well defined. Recently a cardioprotective E2/ERα signaling axis was identified in the pressure overloaded RV---E2/ERα upregulation of apelin, a pro-contractile and pro-survival peptide.^10^ Still, much remains to be elucidated as RV remodeling entails changes in not only contractility but also other determinants of heart function such as cardiac structure and conduction.

In this study, we sought to elucidate mechanisms of sexual dimorphism in the RV structural remodeling response to pressure overload. In patients with advanced HF, RVD is associated with RV interstitial fibrosis.^11^ Hence, we hypothesized that fibrotic remodeling in the pressure-overloaded RV differs by sex and therefore contributes to the sexual dimorphism of RVD. To focus on RV pathophysiology independent of pulmonary vascular pathology, we subjected male and female adult mice to pulmonary artery banding (PAB) for up to 9 weeks. We then serially examined RV structure, function, physiology, and histology. We identified sex-based differences in the onset, extent, and type of cardiac fibrosis that correspond with RV contractile (dys)function. Moreover, we discovered an association between activation of chronic pro-fibrotic inflammation, ECM instability, and maladaptive RV remodeling and failure in males. Our findings suggest that targeting ECM instability and chronic inflammation may be potential approaches for preventing or reversing the deterioration of compensated RVD into RVF.

## METHODS

### Animal use and care

Mice were housed at constant room temperature (23°C), 60±5% relative humidity and fixed 12-hour light/12-hour dark cycle, with *ad libitum* access to food and water. Procedures involving mice conformed to the Guide for the Care and Use of Laboratory Animals by the US National Institute of Health and were approved by the Institutional Animal Care and Use Committee at Columbia University. Animal facilities meet international standards and are regularly checked by certified veterinarians who monitor animal health and welfare and review experimental protocols and procedures.

### Pulmonary artery banding

Male and female *wild-type* mice (C57BL/6: RRID:IMSR_JAX:000664; RRID:MGI:2159965), 10-12 weeks of age, were subjected to or thoracotomy alone (Sham) under surgical plane anesthesia (ketamine, 100mg/kg, i.p., and xylazine, 5mg/kg, i.p.) as previously described.^12^ Briefly, under a dissecting microscope, mice were intubated and connected to a rodent ventilator (Hugo Sachs Elektronik - Harvard Apparatus, D-79232 March, Germany). An incision of the chest was made in the third intercostal space and the pericardium was opened. Following isolation of the pulmonary artery, a suture ligature (Prolene™ 7-0, Ethicon, Puerto Rico, USA) was placed around and tied against a 25-gauge blunted needle which was then rapidly removed. The chest was closed (Polymend® MT, absorbable, 5/0, VPL Sutures, Pensacola, FL, USA), the mouse was extubated, and allowed to recover from anesthesia. All mice received a subcutaneous injection of Meloxicam SR (2mg/mL) for peri-operative and post-surgical analgesia. The severity of RV pressure overload was assessed by echocardiographic measurement of the PA peak pressures. Increased RV pressure was also confirmed by the LVEc. To be included in our study, PAB mice had to have echocardiographic PA peak pressure >10mmHg and LVEc <0.8. Post-surgical mortality was recorded daily. The primary endpoint of the main study was 9wks post-surgery. A survival substudy of PAB mice (6 males and 6 females) followed the animals until their natural death. Survival rate was calculated accordingly.

### Echocardiography

Echocardiography was performed at 3, 6 and 9wks post-surgery in all mice using a digital ultrasound system (Vevo 3100 Image System, Fujifilm, VisualSonics). Additional echocardiographic imaging was performed at 28 and 44wks post-PAB for the survival study subgroup of PAB mice (**Table S1**). Mice were anesthetized with 3% inhaled isoflurane mixed with 1-2L/min O_2_. Vevo LAB 3.0 ultrasound analysis software (Fujifilm, VisualSonics) was used to measure and analyze echocardiographic images. Basal- and mid-RV end diastolic diameter, TAPSE, RV FAC, RV S’, PA peak and mean gradients, and velocity time intervals of flow across the pulmonary valve, tricuspid valve, and pulmonary artery were measured according to the Guidelines and Standards of the American Society of Echocardiography.^13,14^ Off-line analysis was performed by two investigators (I.R., W.D.) blinded to the experimental groups.

### Cardiac magnetic resonance

CMR was performed as previously described^15^ using a horizontal bore 9.4T USR preclinical MRI system (Bruker BioSpin, Germany) with self-gating imaging (IntraGate), and equipped with a linear volume coil and an anatomical shaped surface coil. Cine-MRI pulse-sequences were: matrix 256×128pixels, FOV 25×25mm, echo time 2.9msec, flip angle 15°, slice thickness 1mm, repetition time 12msec, repetitions 1, oversampling 350, movie frames 16, scan time 6’43”200msec. Anesthetized mice (1-3% isoflurane with 1-2L/min O_2_) were pronated in a purpose-built cradle with subcutaneous ECG leads and a rectal probe for ECG, respiration, and body temperature monitoring (SA Instrument, Stony Brook, NY). Following the coronal, sagittal and axial tri-pilot imaging, a horizontal long-axis was obtained to acquire 1mm serial 5-7 short-axis slices covering the entire RV. DICOM images were used for offline analysis of RV volumes, RVEF, RV free wall thickness, end-diastolic diameter, and right atrium area using Image J software (v1.52q). The ratio between RV end-systolic volume and stroke volume was used to assess RV function.^16,17^ The CMR investigator (I.R.) was blinded to the experimental groups during image acquisition and analysis.

### RV pressure-volume (PV) loop recordings

Blinded to the study groups, an investigator (W.D.) performed invasive hemodynamic assessment by cardiac catheterization and PV loop analysis of the RV. RV cardiac catheterization was performed using a closed-chest method whereby the appropriately sized 1F conductance catheter (PVR-1030 or PVR-1035, Millar Instruments) was inserted into the right external jugular vein and advanced into the RV. Upon hemodynamic stability, steady-state baseline data were collected by the conductance catheter coupled to a Millar MPVS Ultra and PowerLab 16/35 data acquisition system (AD Instruments). RV pressure and volume waveforms were recorded simultaneously and analyzed over at least 10 consecutive cardiac cycles. Cuvette calibration of the volume/conductance signal was performed using fresh, heparinized blood (at animal body temperature) and was then followed by saline calibration, as previously described.^18^ HR, SV, CO and RVESP were determined from PV loops. Stroke work (SW) was calculated as SV multiplied by mean pulmonary arterial pressure. A parameter of diastolic function, the relaxation time constant τ was calculated by Weiss method (regression of log (pressure) versus time). Maximal RV power (Powmax), the rate of external work performed, was calculated as SW*HR. Given the differences in body weight (BW) between male and female mice, volumetric parameters were normalized to body weight. All calculations were performed using LabChart 8 software (AD Instruments).

### Immunohistochemistry, histology, and specialized microscopy

Perfusion-fixation of hearts were performed for histological analyses. Mice were euthanized with Euthasol (120mg/kg i.p.), followed by potassium chloride (KCl, 2.5M, i.v., Sigma, #P9541), and then perfused with 4% paraformaldehyde (Fisher Scientific, #AA47377-9M). The heart was excised and then embedded in paraffin. Tissue sections were cut at 5μm thickness.

### Measurement of collagen deposition and replacement and perivascular fibrosis

To examine collagen fibers in the RV, tissue sections at the RV mid-cavity level were dually stained with PR/FG. The amount of collagen in the entire RV cross-sectional area was measured as a percentage of the entire RV. Replacement fibrosis was identified on PR/FG dual stained RV tissue as an area of dead myocardium (defined as cytolytic cardiomyocytes areas) replaced by collagenous scar and characterized by expanded ECM network.^19^ Areas of replacement fibrosis were identified at the mid-myocardial level and expressed as % of the entire RV. Perivascular fibrosis was measured in at least 5 intramyocardial coronary vessels per heart by assessing the area of peri-adventitial collagen. The ratio of periadventitial collagen to the medial area of the vessel was used to quantify perivascular fibrosis.

#### Collagen structure

Two-photon and SHG microscopy were used to assess collagen structure in Sham and PAB mice at 6 and 9wks post-surgery, as previously described.^20^ Images were acquired using an SP8 laser scanning system (Leica Microsystems, Mannheim, Germany), equipped with an HCX APO 20x/0.5 water immersion objective and excited at 830 nm with infrared light produced by a MaiTai Deep See tunable pulsed laser with a pulse width of ∼70fs and an 80MHz repetition rate (Spectra-Physics, Mountain View, CA). Typical laser power was 50%, corresponding to 105mW total power at the sample. The backscattered SHG emission was collected with a HyD detector in the range 385-430nm and the autofluorescence of tissue was collected using a photomultiplier (PMT) detector, in the range 480-525nm. Single-plane images were acquired using the LAS X software (Leica) with a pixel size of 0.541μm. SHG images were analyzed with ImageJ software (v1.52q). Thresholds of images were determined automatically by the Otsu method. The area occupied by SHG signal, and the mean intensity within this area, were calculated.

#### Collagen degradation

To assess collagen degradation within the RV, we used F-CHP (3Helix, Salt Lake City, UT) as previously described.^21^ The amount of RV collagen degradation was assessed in at least 5 fields from one mid-cavity section and expressed as percentage of the F-CHP-positive stained area.

#### Wheat germ agglutinin staining for evaluation of cardiomyocyte size

Cardiomyocyte size in the RV was assessed using WGA staining (1:100 dilution, Alexa Fluor 594 conjugate, Invitrogen, #W11262). The RV cardiomyocyte size was measured for at least 40 cardiomyocytes per heart and the mean RV cardiomyocyte cross-sectional area was calculated.

#### Mac-2 staining for evaluation of myocardial inflammation

To assess RV inflammation, perfusion-fixed, hearts were stained with immunofluorescent rat anti-Mac-2 primary antibody (1:200 dilution, Cedarlane Laboratories, Burlington, Ontario, Canada) followed by secondary antibody incubation (1:200 dilution, goat anti-rat IgG H&L, Alexa Fluor® 488, Abcam, ab 150157,). The density of Mac-2 positive macrophages in the RV was determined by the total cell count in at least 5 random visual fields of the mid RV free wall CSA divided by the total (tissue) surface area of the visual fields. Cell density was expressed as cell/mm^2^. F-CHP-, WGA-, and Mac-2-staining were imaged using a Leica DMi8 Inverted Fluorescent Microscope (Leica Microsystems, Buffalo Grove, IL). Histological quantifications were performed using Image J software (v1.52q) by an investigator (I.R.) blinded to the study groups.

### Transcript analysis by RT-qPCR

Total RNA was extracted from mouse RV tissue using the Tissue RNA Purification Kit (Norgen Biotek) and motorized tissue grinder (Fisher Scientific). Total RNA of each RV sample was quantified using NanoDrop-2000c (Thermo Fisher Scientific). Following total RNA extraction and quantification, cDNA was amplified using primer pairs listed in **Table S2**, SsoFast EvaGreen Supermix, and the CFX96 thermal cycler apparatus (Bio-Rad) as per manufacturer’s instructions. Ribosomal RNA *18S* was used as internal control. RT-qPCR reactions were performed in triplicate. Transcript expression was calculated using the ΔΔCT method.

### Western analysis

Total protein was extracted from RV tissues. Immediately following surgical removal from experimental animals, the RV was quickly frozen in liquid nitrogen. Snap-frozen RV samples were then homogenized with a BeadBug™ Microtube Homogenizer (Alkali Scientific Inc.) in T-PER Tissue Protein Extraction Reagent (Thermo Fisher Scientific) supplemented with a cocktail of 10% protease/phosphatase inhibitors (Sigma Millipore). Tissue homogenates were centrifuged at 10,000g for 5 min at 4°C, supernatant was collected, and protein concentration was measured by standard BCA method. For each sample, 30μg of total protein was subjected to SDS-PAGE (4-20%) under denaturing conditions. Revert™ staining (LI-COR, Lincoln, NE, USA) of transfer membrane was performed to quantify total protein in each lane. Transfer membranes were destained using Revert™ Destaining Solution (LI-COR), then washed clean using Revert™ 700 Wash solution (LI-COR). Western blots were done to detect RV levels of Mac-2 (1:1000 dilution, rat anti-Mac2 antibody, Cedarlane, CL8942AP), Tim-4 (1:1000 dilution, rabbit anti-TIM-4 antibody, Invitrogen, PA5-116045, RRID:AB_2900679) and Lumican (1:1000 dilution, rabbit recombinant anti-Lumican antibody, Abcam, ab168348, RRID:AB_2920864). Secondary antibody incubations were performed using IRDye® 680RD or 800CW as appropriate (LI-COR). Immunoblots were analyzed using the Odyssey® Classic Imaging System (LI-COR Biotechnology, Lincoln, Nebraska, USA).

### Statistical analysis

Statistical analyses were performed using: 2-way ANOVA when determining interaction of conditions or the source of variance, followed by Tukey’s multiple comparison testing, as appropriate; unpaired Student’s t test as appropriate, and with Welch’s correction if variances were unequal. For all analyses, normal distribution was tested using the Shapiro-Wilk normality test. Accordingly, when appropriate, the Mann-Whitney test was applied for nonparametric distributions. Correlation (Spearman r) analysis was used, when appropriate. Survival analysis was performed using the Kaplan-Meier method. Mortality was compared using the log rank test. Fisher’s exact test was used, when appropriate. Statistical significance was defined as p<0.05. GraphPad Prism 10.1.1 was used for statistical and graphical analysis.

## RESULTS

### Despite similar hypertrophic responses to pressure overload, male mice developed RVD sooner and more severely than did female mice

To focus on sex differences in the intrinsic myocardial response of the RV to chronic pressure overload, we subjected both male and female adult mice to PAB for up to 9 weeks. The severity of RV pressure overload was similar between male and female PAB mice (M-PAB, F-PAB) as reflected by their invasively measured RV end-systolic pressure (RVESP) and echocardiography derived LV systolic eccentricity indices (**Table 1**, **Figure S1**). Both sexes had comparable, robust RV hypertrophic responses to PAB as shown by their increased Fulton’s indices and cardiomyocyte cross-sectional area relative to Sham counterparts (**Figure 1B**, **Figure 2**).

**Figure 1.**
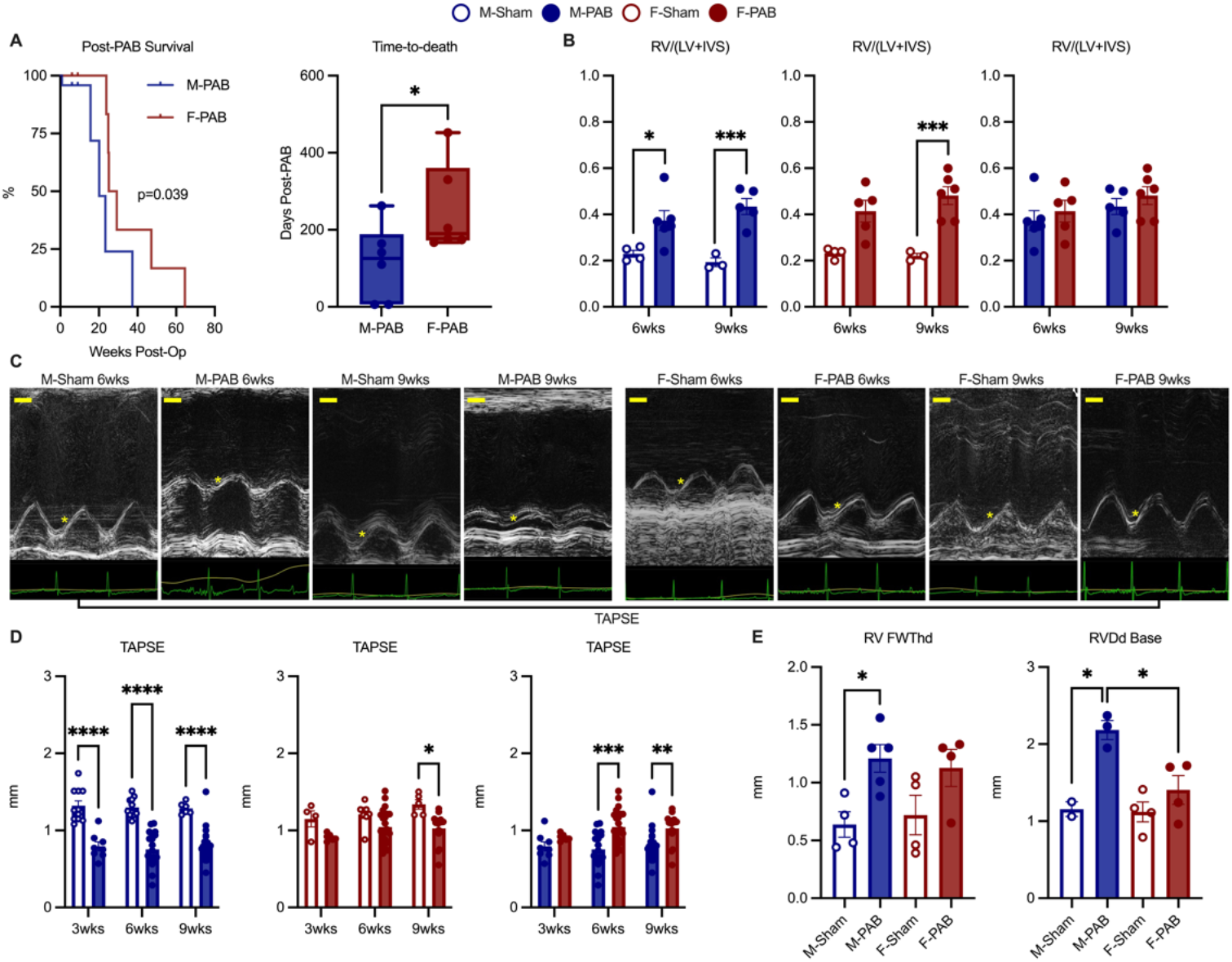
Male mice exhibit worse RV remodeling response to pressure overload than do female mice. **A.** M-PAB had lower survival rate and earlier median time-to-death compared to F-PAB. Kaplan Meier curves for period following PAB. P on Log-rank test. Whisker box plots (median, box: 25th to 75th percentiles, whiskers: min to max). *p=0.026 on Mann Whitney test. **B.** Similar RV hypertrophic responses in M-PAB and F-PAB at both 6 and 9wks post-PAB. Gravimetric parameter of RV hypertrophy shown is Fulton’s index (RV/(LV+IVS)). Bars (mean ± SEM). P values on Student’s t-test. **C.** Representative echocardiographic M-mode images of TAPSE for M-Sham, M-PAB, F-Sham, and F-PAB at 6 and 9wks post-surgery. Yellow * shows TAPSE. Scale bar=1mm. **D.** RV systolic function response to pressure overload was worse in M-PAB than F-PAB. Bars (mean ± SEM). *p<0.05, **p<0.01, ***p<0.001. ****p<0.0001 on 2-way ANOVA followed by Tukey’s test. **E.** CMR analysis of RV Free wall thickness (RV FWThd) and RV basal diameter (RVDd Base) at 6wks post-surgery. Bars indicate mean ± SEM. P values from Student’s t-test. *p<0.05.

**Figure 2.**
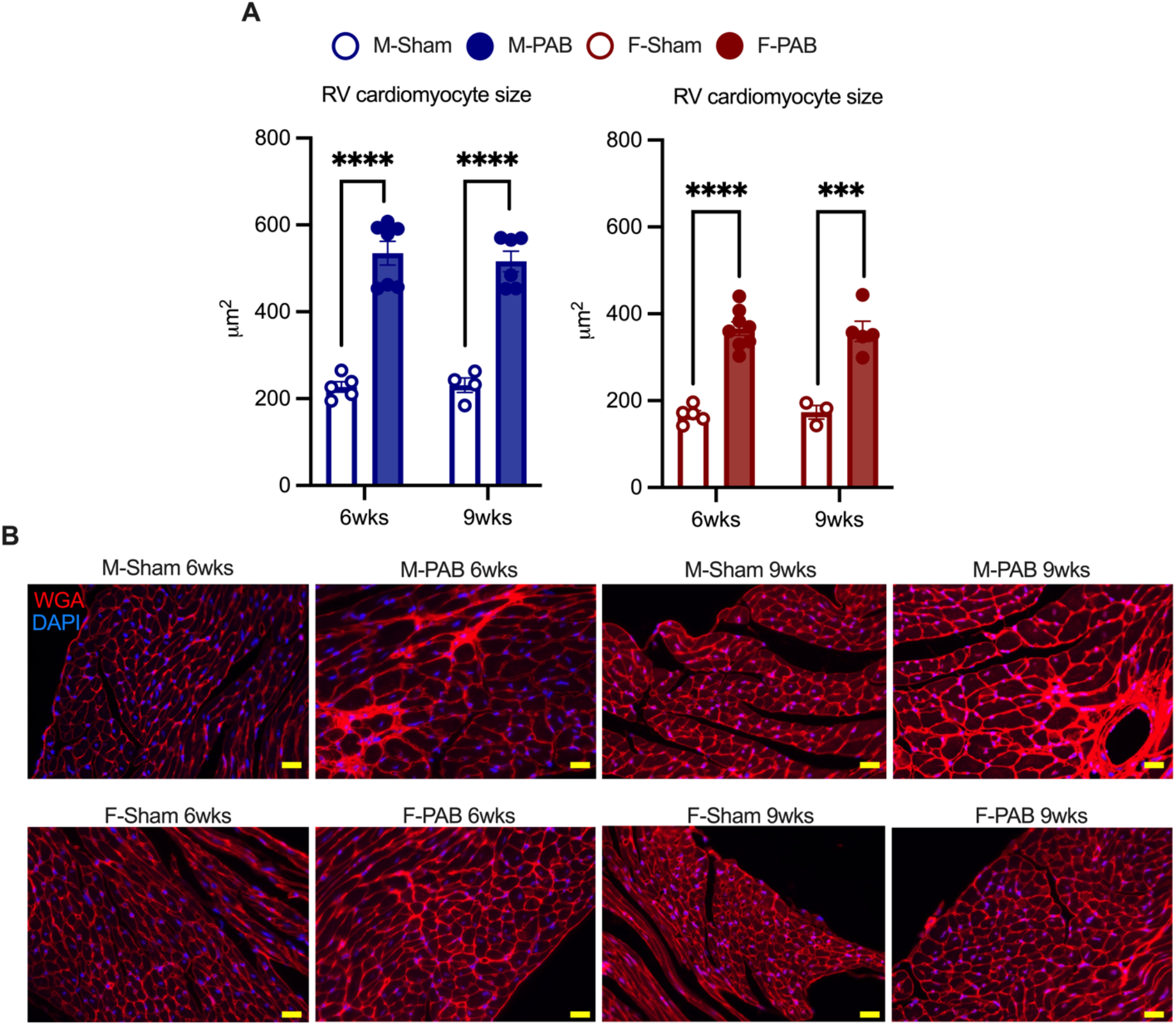
RV cardiomyocyte hypertrophic response to pressure overload is similar between male and female PAB mice. **A.** Cross-sectional area of RV cardiomyocytes increased after PAB in male and female mice compared to sex-corresponding Sham groups. Bars (mean ± SEM). ***p<0.001, ****p<0.0001 on Student’s t-test. **B.** Representative histology images of WGA (red) of RV for Sham and PAB, male and female mice at 6 and 9wks post surgery. DAPI stain, nuclei (blue). Images taken at 400x magnification. Scale bar=25μm.

**Table 1.**
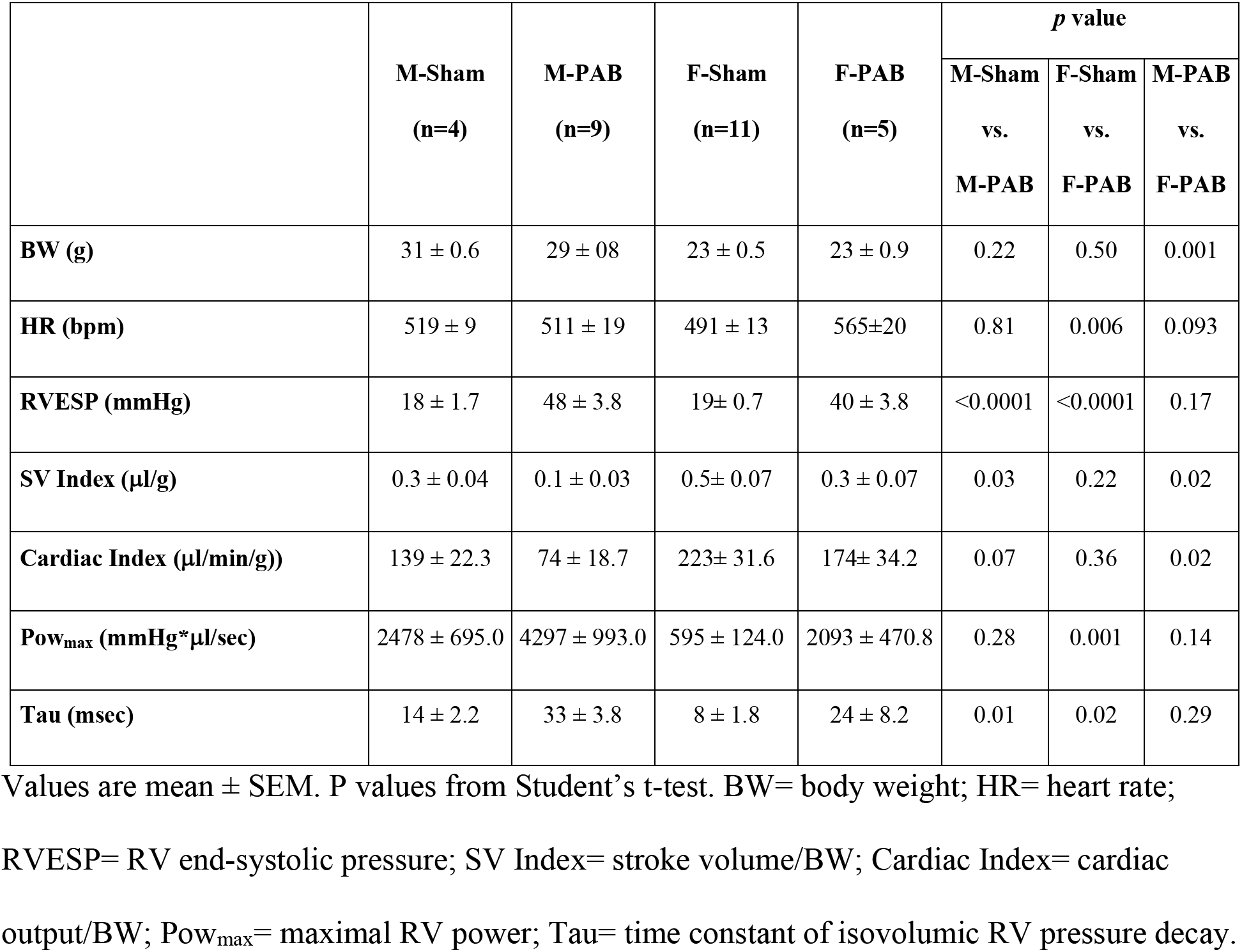
Hemodynamic parameters and indices of RV function at 9wks post-Sham/PAB.

|  | **M-Sham**<br>**(n=4)** | **M-PAB**<br>**(n=9)** | **F-Sham**<br>**(n=11)** | **F-PAB**<br>**(n=5)** | *p* value |  |  |
| --- | --- | --- | --- | --- | --- | --- | --- |
|  |  |  |  |  | **M-Sham**<br>**vs.**<br>**M-PAB** | **F-Sham**<br>**vs.**<br>**F-PAB** | **M-PAB**<br>**vs.**<br>**F-PAB** |
| **BW (g)** | 31 ± 0.6 | 29 ± 0.8 | 23 ± 0.5 | 23 ± 0.9 | 0.22 | 0.50 | 0.001 |
| **HR (bpm)** | 519 ± 9 | 511 ± 19 | 491 ± 13 | 565±20 | 0.81 | 0.006 | 0.093 |
| **RVESP (mmHg)** | 18 ± 1.7 | 48 ± 3.8 | 19± 0.7 | 40 ± 3.8 | <0.0001 | <0.0001 | 0.17 |
| **SV Index (μl/g)** | 0.3 ± 0.04 | 0.1 ± 0.03 | 0.5± 0.07 | 0.3 ± 0.07 | 0.03 | 0.22 | 0.02 |
| **Cardiac Index (μl/min/g))** | 139 ± 22.3 | 74 ± 18.7 | 223± 31.6 | 174± 34.2 | 0.07 | 0.36 | 0.02 |
| **Pow<sub>max</sub> (mmHg\*μl/sec)** | 2478 ± 695.0 | 4297 ± 993.0 | 595 ± 124.0 | 2093 ± 470.8 | 0.28 | 0.001 | 0.14 |
| **Tau (msec)** | 14 ± 2.2 | 33 ± 3.8 | 8 ± 1.8 | 24 ± 8.2 | 0.01 | 0.02 | 0.29 |

Values are mean ± SEM. P values from Student's t-test. BW= body weight; HR= heart rate;
RVESP= RV end-systolic pressure; SV Index= stroke volume/BW; Cardiac Index= cardiac output/BW; Pow<sub>max</sub>= maximal RV power; Tau= time constant of isovolumic RV pressure decay.

Despite equal RV hypertrophy, M-PAB and F-PAB differed in their onset and decrement of RV contractile response. Serial echocardiography (**Figure 1C, D**) detected severe RV systolic dysfunction (RVD) in M-PAB at 6wks post-surgery (TAPSE, M-Sham6wks vs. M-PAB6wks: 1.3±0.04 vs. 0.8±0.05mm, p<0.0001) which persisted unchanged to 9wks post-surgery (TAPSE, M-PAB6wks vs M-PAB9wks: 0.8±0.05 vs. 0.8±0.04mm, p=0.81; M-Sham9wks vs. M-PAB9wks: 1.3±0.03 vs. 0.8±0.04, p<0.0001). In contrast, F-PAB6wks maintained RV function comparable to that of F-Sham6wks (TAPSE, F-Sham6wks vs. F-PAB6wks: 1.2±0.05 vs. 1.0±0.05mm, p=0.31). Hence, M-PAB6wks developed RVD while F-PAB6wks maintained normal RV systolic function (p<0.001). By 9wks post-surgery, F-PAB RV function did decline but only mildly (TAPSE, F-Sham9wks vs. F-PAB9wks: 1.3±0.06 vs. 1.0±0.05mm, p=0.024). Accordingly, RVD was less severe in F-PAB9wks than in M-PAB9wks (p=0.009).

Subsequent CMR and RV PV loop analyses substantiated the sex differences that we had detected by echocardiography. At 6wks post-surgery, M-PAB showed increased RV free wall thickness (RV FWThd) and greater RV basal dilation (RVDd-base) compared to M-Sham (**Figure 1E, Videos S1-S4**). F-PAB demonstrated neither RV free wall thickening nor RV dilation. In fact, RV FWThd and RVDd-base were similar for F-PAB6wks and F-Sham6wks. By 9wks post-surgery, M-PAB developed RVF as revealed by reduction of both RV stroke volume (RV SV) and RV cardiac output (RV CO) (**Table 1**). F-PAB9wk did not develop RVF despite the presence of RVD; there was no difference in RV SV and RV CO between F-PAB and F-Sham (**Table 1**). The sex differences in RV structural and functional remodeling in PAB mice mirrored that of PAH patients and HF patients with pHTN; men respond with maladaptive RV remodeling while women respond with adaptive (compensatory) RV remodeling. Consistent with these patterns of RV remodeling, F-PAB had a higher survival rate (log-rank HR 2.9, 95% CI 0.8-10.4, p=0.039) and longer survival duration than M-PAB (18 [0.8-27] vs. 25 [24-38]) wks, p=0.026, **Figure 1A**).

### Extracellular matrix remodeling of pressure-overloaded RV differs between male and female mice

In HF patients, cardiac fibrosis has been shown to be inversely associated with ventricular function.^22^ Hence, we sought to identify sex differences in the RV extracellular matrix (ECM) remodeling response to pressure overload. Total collagen was increased in the RV of both M-PAB and F-PAB, relative to their respective Sham (**Figure 3A**). However, the temporal dynamics and amount of collagen deposition in the RV differed between M-PAB and F-PAB. Collagen deposition increased throughout the 9wk post-surgical period for M-PAB but appeared to plateau at 6wks post-surgery for F-PAB. Additionally, the increase in RV collagen deposition and fibrosis was greater and more extensive in M-PAB than in F-PAB. (**Figure 3A)**. The pattern of fibrosis in M-PAB was predominantly diffuse and interstitial, with sparse focal areas of replacement fibrosis throughout the RV (**Figure 3B, C**). We also analyzed perivascular fibrosis (PVF, ratio of periadventitial collagen to medial area of the vessel). At 6wks post-surgery, PVF in the RV was greater in M-PAB compared to M-Sham (p=0.010) but not so in F-PAB vs F-Sham (**Table S3**). Similarly, RV PVF increased further, though non-significantly, in M-PAB9wks (p=0.22). No difference in PVF was noted between F-PAB9wks and F-Sham9wks (p=0.94). By 9wks post-surgery, PVF of M-PAB was non-significantly higher than that of F-PAB9wks (p=0.18). Differential expression of genes related to fibrosis and ECM remodeling also suggested male-specific activation of pro-fibrotic pathways and other sex-dependent ECM remodeling programs in the pressure overloaded RV (**Figure S2**).

**Figure 3.**
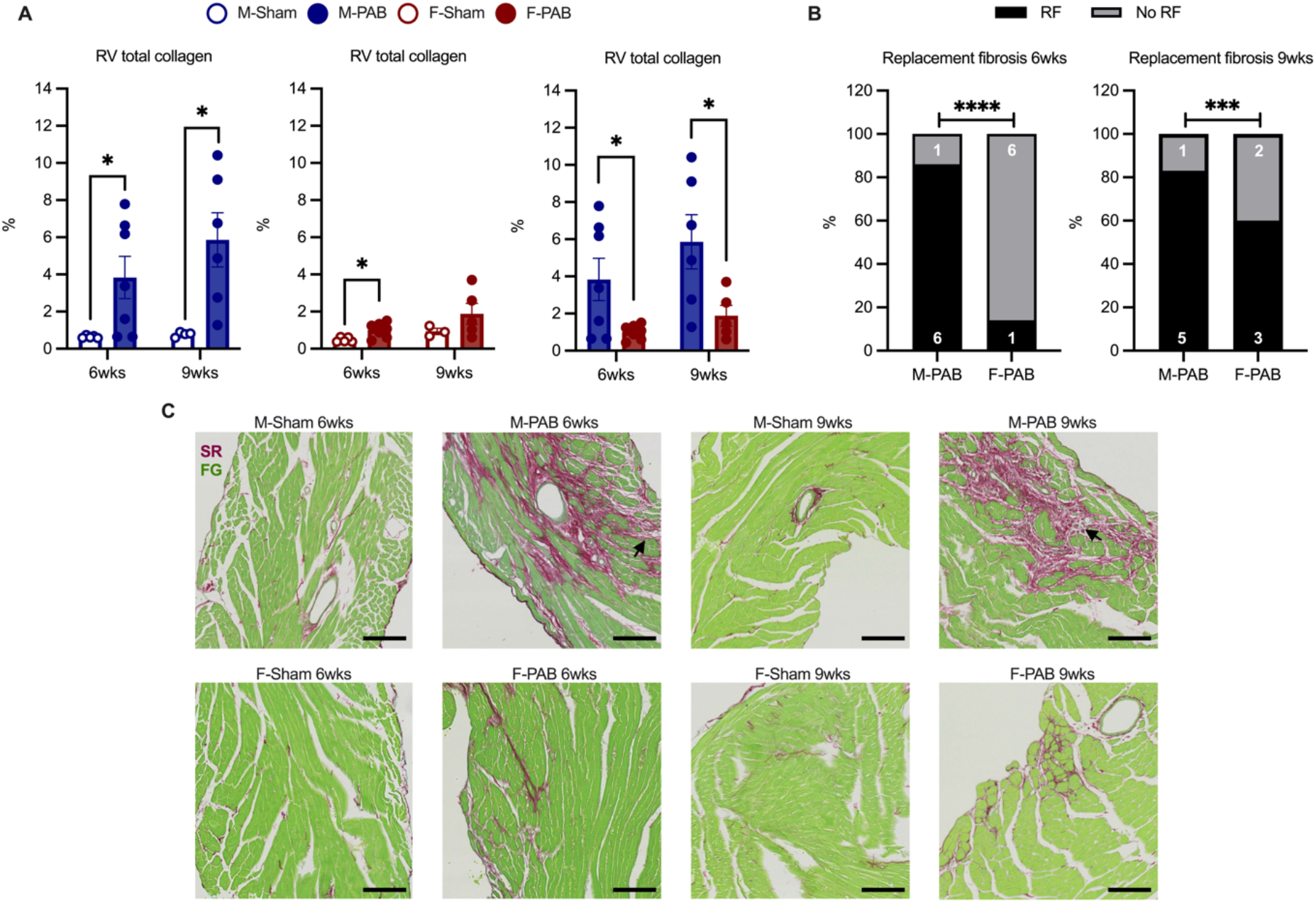
RV fibrotic remodeling response to pressure overload was more severe in male mice. **A.** Quantitative analysis of RV total collagen for male and female mice (Sham and PAB). Bars (mean ± SEM). *p<0.05 on Student’s t-test. **B.** Chi square analysis of RV replacement fibrosis (RF) in PAB mice at 6 and 9wks post-PAB. Bars (% of mice). Numbers indicate the number of mice within the designated subset. ***p<0.001, ****p<0.0001 on Fisher’s test. **C.** Histology images of the RV showing collagen deposition in M-Sham, M-PAB, F-Sham, and F-PAB mice. Black arrows indicate areas of replacement fibrosis. Images taken at 200x magnification. Scale bar=100μm.

ECM remodeling is a dynamic process that integrates collagen deposition with collagen degradation. Thus, we subsequently analyzed collagen degradation in the RV via Collagen Hybridizing Peptide 5-FAM conjugate (F-CHP) labeling. Fluorescent staining with CHP revealed an increased pattern of collagen degradation in M-PAB but not F-PAB, relative to their respective Sham (**Figure 4A, B**). The temporal dynamics of collagen degradation was also like that of collagen deposition in M-PAB; collagen degradation increased in M-PAB, compared to M-Sham, at both 6 and 9wks post-surgery. In contrast, RV collagen degradation was no different between F-PAB and F-Sham at either time points. Moreover, RV collagen degradation at 6wks post-surgery was greater in M-PAB than F-PAB (F-PAB6wk vs M-PAB6wk, 2±0.8 vs. 8±2.3%, p=0.036). By 9wks post-surgery, the extent of RV collagen degradation was still greater, albeit non-significantly, in M-PAB9wks than in F-PAB9wks (3±1.0% vs. 7±2.3%, p=0.14). These findings suggest that early and pronounced RV collagen degradation distinguishes M-PAB from F-PAB.

**Figure 4.**
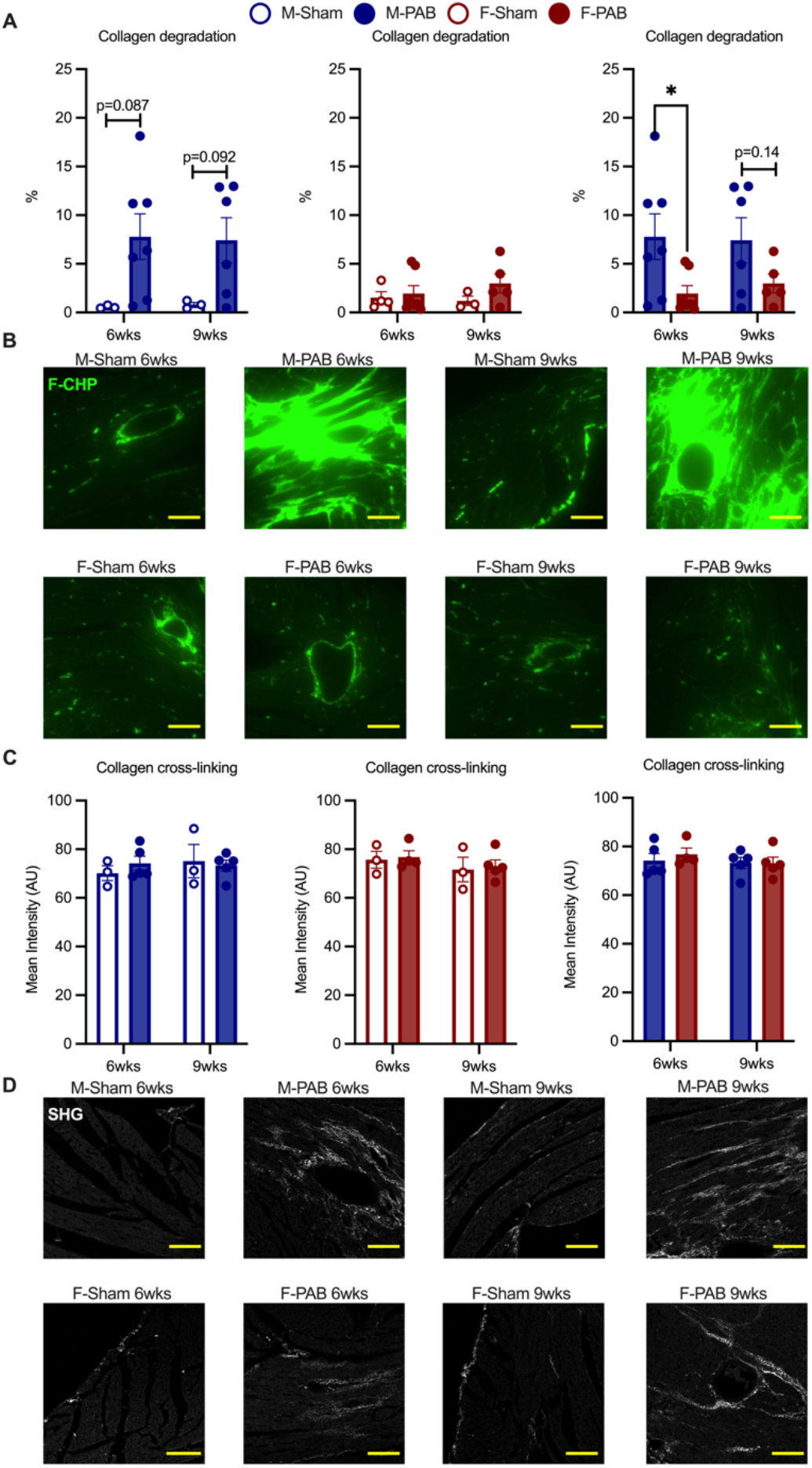
Pressure overload induced collagen degradation without affecting collagen cross-linking in male mice. **A.** Quantitative analysis of degraded collagen in the RV of Sham and PAB mice using F-CHP staining. Bars represent mean ± SEM. *p<0.05 from Student’s t-test. **B.** Representative images of F-CHP staining (green) are shown. Magnification at 200x. Scale bar=100μm. **C-D.** Quantitative analysis (Student’s t-test) and representative images of collagen cross-linking by SHG signal (bright dots) produced by Sham and PAB mice. Bars (mean ± SEM). Images taken at 200x. Scale bar=100μm.

We then sought to determine the effect of the observed differential ECM remodeling dynamics on collagen structure. We measured collagen fiber thickness using two-photon and second-harmonic generation (SHG) microscopy. SHG mean intensities were no different between PAB and Sham counterparts in either sex, indicating that RV collagen fiber thickness was not affected by pressure overload; there was no increased collagen cross-linking with PAB (**Figure 4C, D**). In M-PAB, the earlier onset of fibrosis and collagen degradation in the RV paralleled the time course of RVD development. Taken together, these findings suggest that ECM remodeling of the pressure overloaded RV differs by sex, with males exhibiting a worse phenotype.

### Inflammatory response in the RV is profound in male but not female PAB mice

Inflammation plays a central role in ECM remodeling, and these two processes often act in a reciprocal, positive feedback nature. For example, some metalloproteinases (e.g., MMP2 and MMP9) not only degrade ECM components but also regulate pro-inflammatory cytokines, thereby modulating inflammation.^23^ Thus, we sought to determine whether the pressure-overloaded RV exhibited any sex differences with regards to inflammatory activation and, if so, whether those sex differences were associated with those we found in RV ECM remodeling.

To examine chronic inflammation in the RV, we quantified the density and absolute number of Mac-2+ activated macrophages, mediators of chronic inflammation. We observed increased macrophage density in M-PAB at both 6 and 9wks post-surgery (M-Sham6wks vs. M-PAB6wks, 19±5.4 vs. 47±11.3 cells/mm^2^, p=0.17; M-Sham9wks vs. M-PAB9wks, 4±1.5 vs. 35±9.3 cells/mm^2^, p=0.030, **Figure 5A, C**). No such increase was observed in F-PAB RV, at either time points (**Figure 5A, C**). In addition, the absolute number of macrophages in the RV was consistently higher in M-PAB than in time-matched F-PAB (**Figure 5A, C**). As activated macrophage density in the RV increased, so did collagen degradation (**Figure 5B**). These histological findings were corroborated by Western analysis of MAC-2 protein in the RV. At 6wks post-PAB, RV MAC-2 expression increased by 52% in M-PAB compared to that of F-PAB (p=0.029). RV expression levels of TIM-4, a marker of cardiac resident macrophages, was comparable between M-PAB and F-PAB. These patterns in MAC-2 and TIM-4 expression suggest that the number of activated macrophages in the RV increased in M-PAB but not F-PAB (**Figure 5D, E**).

**Figure 5.**
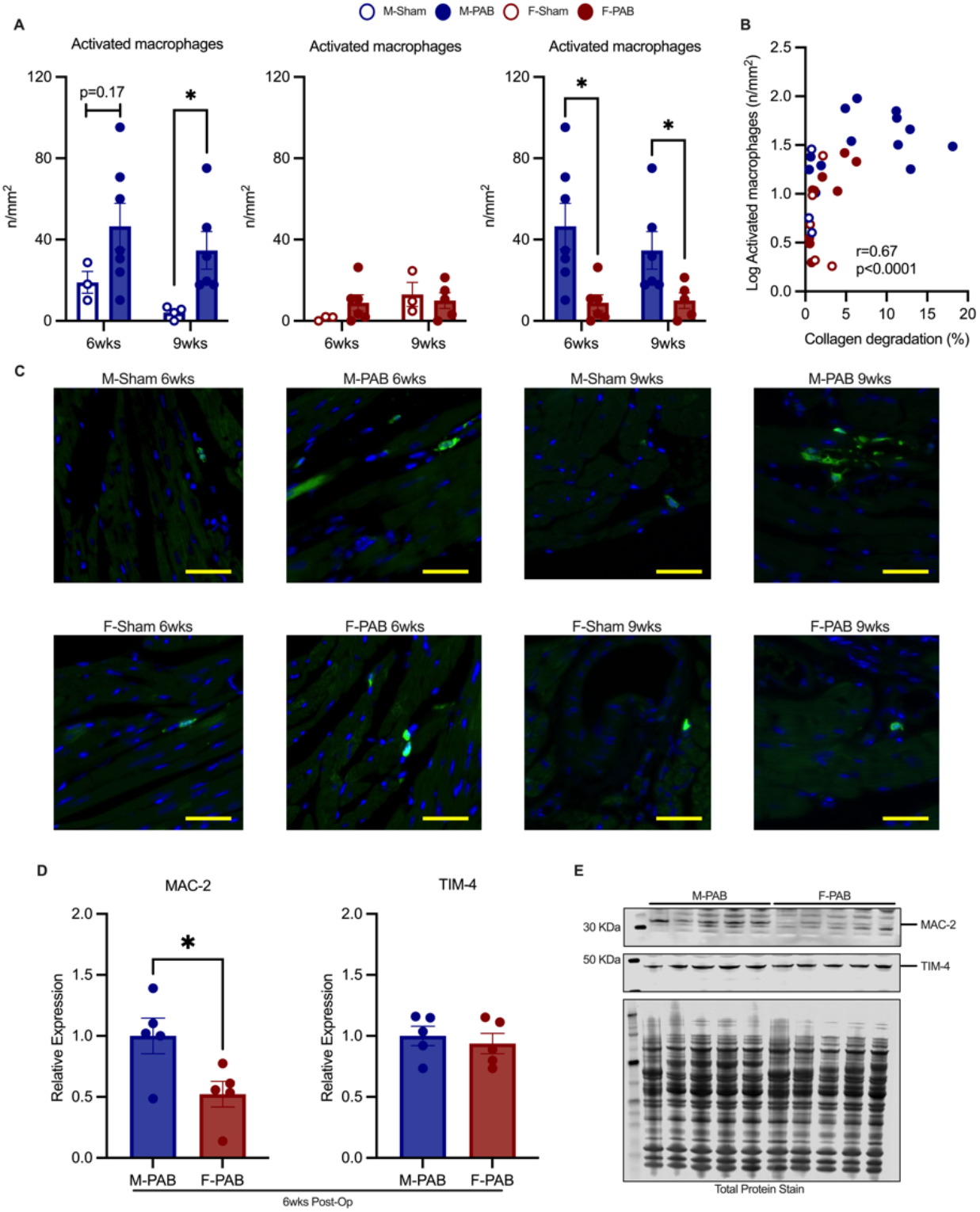
Increased inflammatory response in male mice after PAB. **A.** Macrophages quantification in the RV using immunofluorescence staining with Mac-2. Assessment performed in Sham and PAB male and female mice at 6 and 9wks after PAB. Mean ± SEM shown. *p<0.05 from Student’s t-test. **B.** Positive correlation between collagen degradation (F-CHP) and macrophages number in the RV. Spearman r correlation coefficient and p value shown. **C.** Immunofluorescence images of Mac-2 staining for macrophages (green) in M-Sham, M-PAB, F-Sham and F-PAB at 6 and 9wks. Nuclear staining (blue) using DAPI. Images taken at 400x. Scale bar=63.7μm. **D.** Summary densitometry analysis of Western normalized to total protein stain, relative to M-PAB. Bars indicate mean ± SEM. *p<0.05 from Student’s t-test. **E.** Representative Western blots of MAC-2, TIM-4 and total protein stain.

### Extracellular proteoglycan Lumican is deposited in the pressure overloaded RV of female but not male PAB mice

Given our sex-specific findings of ECM remodeling and non-resident macrophage activation in M-PAB RV, we also evaluated potential modulators of ECM stability in the pressure overloaded RVs. We focused on Lumican, a small leucine-rich proteoglycan that regulates collagen structure in the heart, rendering the ECM more stable.^24, 25^ We measured the transcript and protein expression levels of Lumican (*Lum*, LUM) in the RV at 9wks post-PAB. Despite similar *Lum* transcript expression in both sexes (**Figure 6A**), RV expression of LUM protein differed between F-PAB and M-PAB. LUM levels in the RV were higher in F-PAB compared to F-Sham (p<0.001, **Figure 6B, C**). In contrast, this upregulation of LUM in the RV did not occur in M-PAB. Moreover, LUM expression was consistently lower in M-PAB than in F-PAB mice (p=0.001, **Figure 6B, C**). Accordingly, the higher basal expression of LUM and its pressure overload-induced upregulation in F-PAB could help fortify ECM stability. This advantage of an ECM stabilizing response to RV pressure overload may be one mechanism that protects females against maladaptive RV remodeling. Hence, a differential balance of negative and positive ECM regulators, including matricellular proteins and extracellular proteoglycans, likely contribute to the sex differences in RV remodeling phenotypes.

**Figure 6.**
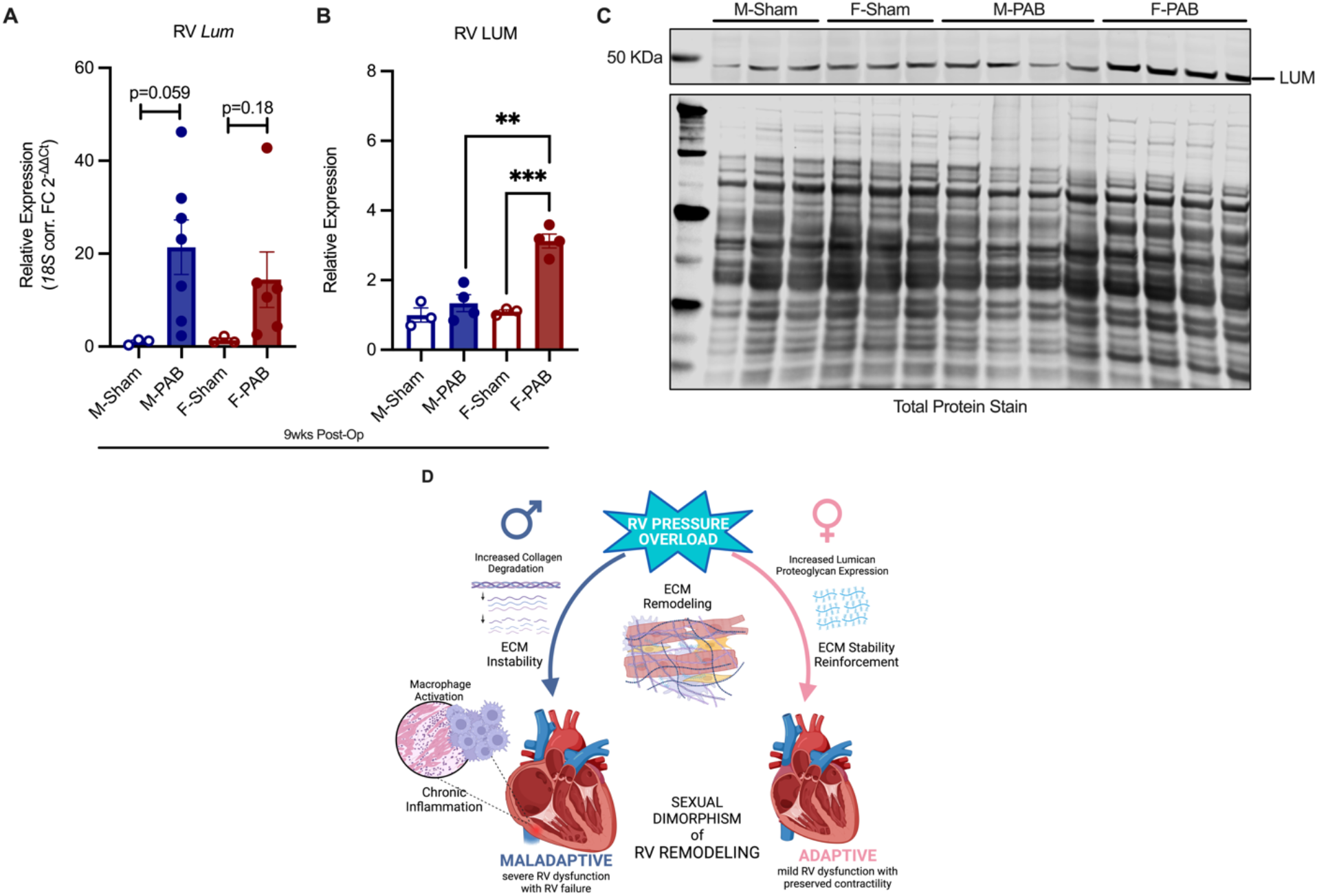
Differential deposition of Lumican may contribute to sex differences in RVD ECM remodeling. RNA and proteins were extracted from the RV of male and female mice subjected to Sham or PAB procedures. **A.** Transcript expression of *Lum* in the RV of Sham and PAB mice at 9wks post-surgery. Bars represent mean ± SEM. P values from Student’s t-test. **C.** Summary densitometry analysis of Western normalized to total protein, relative to M-Sham. Bars indicate mean ± SEM. **p<0.01, ***p<0.0001 from Student’s t-test. **C.** Representative Western blots of LUM and total protein stain. **D.** Our study showed that over 9wks of equal RV pressure overload, male PAB mice developed severe RVD and RVF, while female PAB mice developed only mild RVD. RV ECM instability and chronic RV inflammation distinguished male from female PAB mice. Created with BioRender.com.

## DISCUSSION

Our study is the first to demonstrate a male-specific association between ECM destabilization, chronic inflammatory activation by recruited macrophages, and deterioration of the dysfunctional pressure-overloaded RV into RV failure (**Figure 6D**). No other study, to the best of our knowledge, has investigated both fibrosis and inflammation in this context of pressure overload-induced RV remodeling. Here we identified two novel sex-specific aspects of RV fibrotic remodeling that differentiate decompensated (maladaptive) from compensated (adaptive) RV dysfunction.

### Sex-specific ECM remodeling response to RV pressure overload

By deeply characterizing ECM remodeling, we discovered that RV pressure overload induced collagen degradation in the RV of male but not female mice. Such granular assessment of cardiac ECM remodeling had not been previously performed in any model of RVD or pHTN. Our identification of this select sex-specific ECM remodeling response to RV pressure overload was significant for its impact on ventricular function. The male-specific pressure overload-induced collagen degradation destabilized the RV ECM and thereby contributed to the maladaptive RV remodeling in males. Another distinct finding of our study, with regards to pressure overload-induced RV fibrotic remodeling, is that expression of ECM stabilizer Lumican was upregulated in the RV of female but not male PAB mice. Hence, only female PAB mice benefitted from the cardioprotective action of Lumican in the pressure-overloaded RV.

### Sex-specific chronic inflammation of pressure-overloaded RV

Another salient and novel discovery of our study is the male-specific, chronic activation of recruited macrophages detected in the pressure-overloaded RV. By directly examining cardiac inflammation by histology and molecular analysis of myocardial tissue, we demonstrated that activated macrophages were recruited in the pressure-overloaded RV of male but not female PAB mice. This form of chronic cardiac inflammation was specific to RV pressure overload remodeling in male PAB mice. Given our focus on sex differences, our study was powered to enable comparison of male and female animal models. Rare in numbers, prior clinical and pre-clinical studies of sex differences in RV remodeling had small study cohorts and more limited methodology. In one such clinical study, blood levels of a biomarker of macrophage activation were found to be elevated in PAH patients.^26^ However, without any direct histological evidence of cardiac inflammation, the study findings could be interpreted as chronic inflammation of either the pulmonary vasculature or the heart. Additionally, its small number of study subjects precluded sex-based comparison. To the best of our knowledge, there have not been any animal studies of sex-differences in myocardial inflammation in the pressure overloaded RV. To date, pressure overload-induced myocardial inflammation has been studied in a mouse model of LV pressure overload (transverse aortic constriction, TAC), but not RV pressure overload. Although the study included both sexes, its analysis was limited to pooled data.^21^

### Sexual dimorphism of RV functional remodeling and RVF in PAB mice

By establishing our study endpoint at 9wks post-surgery, we were able to capture, with hemodynamic evidence, the transition from severe RVD to frank RVF in male PAB mice. Likewise, our RV PV loop data showed that even when the female PAB mice developed RVD, they did not have RVF. Other rodent PAB studies typically had shorter post-surgical periods. Despite developing RV hypertrophy and RVD, these rodents with short duration of RV pressure overload did not transition to RVF (concomitant RV dilatation, elevated RVEDP, and diminished RV SV). Hence, these prior rodent PAB studies would not have been able to detect sex differences in the crucial transition from RVD to RVF.

These sex-based differences in pressure overload RV functional remodeling, hemodynamic deterioration to RVF, and death in PAB mice mirror those observed in HF with pHTN and PAH patients. Analogous sex-differences were detected in the abundance and density of activated macrophages recruited in the RV, the time course and severity of collagen deposition and degradation, and the development of ECM instability in the RV. These detrimental histopathologic findings correlated with more severe RVD and ultimately RVF in M-PAB. In contrast, F-PAB did not share these histopathologic findings of M-PAB. F-PAB maintained preserved RV function for a longer period and developed more stable, mild RVD, without RVF. Thus, our findings suggest that sex differences in ECM remodeling and chronic inflammation are important pathophysiological factors that distinguish pressure overload-induced maladaptive RV remodeling in males from adaptive, compensatory RV remodeling in females.

A hallmark of pathologic remodeling, cardiac fibrosis occurs in various heart diseases and in different patterns.^27^ Despite being extensively studied in the LV, cardiac fibrosis and its pathophysiologic role in the pressure-overloaded RV has not been as well characterized. In the present study, we performed a comprehensive analysis of fibrotic remodeling of the pressure-overloaded RV, highlighting sex-based responses in ECM remodeling and patterns of fibrosis. Maladaptively remodeled, M-PAB RV demonstrated activation of a pro-fibrotic gene program, diffuse interstitial fibrosis with few and sporadic small foci of replacement fibrosis, and perivascular fibrosis. In contrast, the adaptively remodeled F-PAB RV showed suppression of pro-fibrotic factors, only a small amount of collagen deposition, no significant perivascular fibrosis, and increased levels of the ECM stabilizing proteoglycan Lumican.

Direct comparison of cardiac fibrosis between both sexes, as we did in this study, has been uncommon. However, our findings are in line with those of chronic diseases of various organs. Female animal models of chronic lung, kidney, and liver diseases have also suggested that estrogen modulates disease progression through anti-fibrotic actions. In these studies, ovariectomized adult female rodents developed worse disease progression, with more severe fibrosis and increased mortality, than those with intact ovaries.^28^ Other studies of female rodents with and without ovariectomy have shown an estrogen-dependent anti-fibrotic signaling pathway in the pressure overloaded ventricle, ^4, 6, 29, 30^ though mechanistic details remain to be elucidated. Aside from sex hormone regulation of fibrosis, sex differences in cardiac fibroblasts (CFs) have also been reported. In a rat model of cardiac fibrosis induced by chronic β-adrenergic stimulation, males exhibited a worse fibrotic phenotype than females; male CFs were more activated than female CFs.^31^

### Sex hormone-dependent vs. -independent modulation of sex specific RV remodeling

Little is known about cardiac inflammatory activation by RV pressure overload, let alone associated sex differences. However, a few studies over the past five years have raised the possibility that modulators other than sex hormones may also underlie the sexual dimorphism of RV remodeling. In a rat PAB model, ovary-intact and ovariectomized females had similar amounts of collagen content in the RV,^29^ indicating that regulation of cardiac collagen content does not depend upon estrogen. Additionally, some sex chromosome-specific genes are associated with pathologic cardiac remodeling but not sex-hormone signaling.^32^ Located in the Y but not X chromosome, several of these genes are involved in the immune response and macrophage maturation. Recent studies have identified dysfunctional regulatory T cells (Tregs)^33, 34^ as significant agents of inflammation in PAH, even suggesting that abnormal Tregs might contribute to sex-disparities in PAH phenotypes.^35^ Their role in RV remodeling, if any, is unknown. Hence, it is possible that our finding of activated non-resident, thus recruited macrophages in the RV of male but not female PAB mice may be due to sex-specific factors that are independent of sex hormones.

### Sex-specific ECM remodeling is associated with sex-specific immune response in maladaptive vs adaptive RV remodeling

We identified an association between pathological activation of macrophages and increased collagen turnover in the maladapted RV of male mice. In the pressure-overloaded RV of M-PAB mice, increased collagen degradation was associated with increased Mac-2+ macrophage density, reflecting intense, chronic inflammatory activation. ECM remodeling in the RV of M-PAB yielded a predominance of interstitial fibrosis. (The observed foci of replacement fibrosis in the RV of M-PAB mice were minimal compared to that reported in LV pressure overload^21^ and myocardial infarct animal models.^36^) These histopathologic findings in the RV of M-PAB mice were absent from F-PAB RVs. Additionally, M-PAB RV did not exhibit the pressure overload-induced upregulation of Lumican that was detected in F-PAB RV. The profound collagen degradation and unchanged low expression of Lumican in M-PAB RV could be due to dysregulation of ECM-preserving pathways, with subsequent activation of proinflammatory cascades. While more studies need to be done to confirm such a sex-specific mechanism, findings in an LV pressure overload model do support a pathogenic role for proinflammatory ECM remodeling. In a fibroblast-specific knockout of TGF-β signaling mediator Smad3, generation of proinflammatory ECM fragments was associated with more severe, pressure overload-induced adverse LV remodeling and deterioration of LV contractile function.^21^ Whether any sex differences existed in that pathophysiologic pathway, however, could not be discerned since the fibroblast-specific Smad3 knockout study pooled data from both male and female mice for analysis. Nonetheless, one could theorize that dysregulation of anti-inflammatory modulators could trigger and promote chronic inflammation in the RV of M-PAB mice.

Conversely, the adaptive RV remodeling of F-PAB could be due to tight regulation of the ECM, expression of specialized cardioprotective matrix proteins and proteoglycans, preserved pro-survival signaling, and induction of anti-fibrotic, anti-inflammatory pathways. Although data are conflicting, sex-related disparities in MMPs levels have been observed in several cardiovascular and neurological disorders.^37^ Whether sex differences in cardiac inflammation and ECM remodeling are driven by sex chromosome-specific genes, sex hormone-regulated signaling, or both have yet to be elucidated.

### Limitations

The present study is admittedly limited. Without genetically modified mice, we cannot interrogate specific pathways of ECM remodeling (i.e., collagen degradation, Lumican cardioprotection) or chronic macrophage activation and their centrality to sex differences in pressure overload RV remodeling. However, our comprehensive histologic, physiologic, and functional comparison of male vs. female wild type PAB mice did identify, in an unbiased manner, likely mediators of the sexual dimorphism in pressure overload RV remodeling, thereby providing direction for new mechanistic studies. Secondly, our molecular analysis of myocardial tissue consisted of RT-qPCR and immunoblotting. Other unbiased methods such as genomic, transcriptomic, or proteomic analyses, or single cell analysis, would have yielded more abundant and granular data. Such big data analyses, however, were beyond the intended scale and scope of our study.

## CONCLUSION

In summary, our study discovered an association between increased collagen degradation, ECM instability, and chronic inflammatory activation by recruited macrophages as pathogenic features underlying, and specific to, the early development of severe RVD and transition to RVF in male PAB mice. Our findings suggest that deeper understanding of sexual dimorphisms in ECM stabilization, fibrosis, and chronic inflammation may potentially offer insights into delaying, or even preventing, RVF in HF and PH patients.

## ACKNOWLEDGMENTS

The authors thank Qing Li for assistance with mouse surgery, and Yanping Sun for guidance with CMR. Analysis of the SHG images was developed in collaboration with Emilia Laura Munteanu from the Confocal and Specialized Microscopy Shared Resource, Herbert Irving Comprehensive Cancer Center (HICCC) at Columbia University Irving Medical Center (CUIMC), supported by the NIH National Cancer Institute (P30CA013696).

## Supplemental Material

Tables S1-S3

Figures S1-S2

Videos S1-S4

## Nonstandard Abbreviations and Acronyms

CMR: cardiac magnetic resonance
ECM: extracellular matrix
F-CHP: collagen hybridizing peptide 5-FAM conjugate
HF: heart failure
LV: left ventricle or left ventricular
LVEc: end-systolic left ventricular eccentricity index
PA: pulmonary artery
PAB: pulmonary artery binding
PAH: pulmonary arterial hypertension
PH: pulmonary hypertension
RV: right ventricle or right ventricular
RVD: right ventricular dysfunction
RVF: right ventricular failure
SHG: second harmonic generation
SR/FG: picrosirius red/fast green

